# Insights in neuronal tuning: Navigating the statistical challenges of autocorrelation and missing variables

**DOI:** 10.1101/2023.10.25.563994

**Authors:** Fredrik Nevjen, Benjamin Adric Dunn

## Abstract

Recent advances in neuroscience have improved our ability to investigate neural activity by making it possible to measure vast amounts of neurons and behavioral variables, and explore the underlying mechanisms that connect them. However, comprehensively understanding neuronal tuning poses challenges due to statistical issues such as temporal autocorrelation and missing variables, as neurons are likely driven in part by unknown factors. The field consequently needs a systematic approach to address these challenges. This study compares various methods for covariate selection using both simulated data and calcium data from the medial entorhinal cortex. We conclude that a combination of cross-validation and a cyclical shift permutation test yields higher test power than other evaluated methods while maintaining proper error rate control, albeit at a higher computational cost. This research sheds light on the quest for a systematic understanding of neuronal tuning and provides insight into covariate selection in the presence of statistical complexities.

## Introduction

Single neuron tuning has been an important tool in our attempts to understand the brain. Many insights have come from considering, e.g., the possible roles of head direction cells (***Taube et al., 1990***), place cells (***O’Keefe and Dostrovsky, 1971***), and grid cells (***Hafting et al., 2005***) – each of which was first characterized through careful inspection of their response properties of single neurons to aspects of the animal’s behavior. As neuroscience moves towards more population-based approaches (***Cunningham and Yu, 2014; Bassett and Sporns, 2017; Datta et al., 2019***), the classification of neurons is still just as important both to leverage traditional methods but also to isolate the relevant cell populations to further study with population-based methods such as dimensionality reduction (***Cunningham and Yu, 2014***), neural manifolds (***Gallego et al., 2017***), latent variable models (LVM) (***Lawrence, 2003; Yu et al., 2008; Wu et al., 2017; Pandarinath et al., 2018; Bjerke et al., 2023; Schneider et al., 2023***), and topological data analysis (TDA) (***Curto and Itskov, 2008; Rybakken et al., 2019; Gardner et al., 2022***). Furthermore, at the same time that technology has enabled large neural recordings, our ability to capture detailed descriptions of animals has also improved (***Mathis et al., 2018***), sparking a renewed interest in animal behavior, while dramatically expanding the number of variables considered to explain the neural activity (***Mimica et al., 2023***). This expansion in terms of neurons as well as possible explanatory variables makes it both more difficult to appropriately ask the question of what variables appear to matter and how we can interpret the answer we receive.

Working under the hope and assumption that neurons have some form of specialization, the need for covariate selection is clear: out of a large number of possible covariates, such as position in space, speed, and head direction, to which features does a neuron selectively respond? This is, however, a deceptively difficult question to ask in a meaningful way, and we wish to highlight important statistical challenges that commonly occur in this endeavor. At their core these challenges boil down to misspecification of statistical models or tests, causing them to yield misleading results. We focus on the generalized linear model (GLM) (***McCullagh and Nelder, 1989***), which is a versatile and popular framework to model the relationship between neural activity and external covariates (***Truccolo et al., 2005; Weber and Pillow, 2017; Hardcastle et al., 2017***), but the issues studied here are important regardless of model choice.

The first challenge we would like to highlight is related to the temporal autocorrelation in neural data. Neural activity and behavioral variables have a non-zero correlation between the values at time t and those preceding and following that time point, arising due to the time-dependent nature of the underlying factors driving the activity, including internal chemical and electrical processes, dynamics of animal behavior, and other external stimuli (***Wilson, 1999; Keat et al., 2001; Truccolo et al., 2005; York et al., 2020; Datta et al., 2019; Hubel and Wiesel, 1959***). Apparent correlations between two autocorrelated time series that appear significant when they are actually independent of each other, known as *nonsense correlation*, were as relevant 100 years ago (***Yule, 1926***) as they are today (***Harris, 2020a***). Figures 1 A and B illustrate how this can influence hypothesis tests concerning correlations if one applies a test assuming independent data. This problem naturally extends to the incorrect use of any statistical method or model that assumes temporal independence. The GLM for instance assumes that the response is independent when conditioned on the variables included in the model (analogous to the error terms in a linear regression model being independent), which will only hold in the unlikely case that the included variables capture all of the underlying temporal structure (autocorrelation).

**Figure 1.**
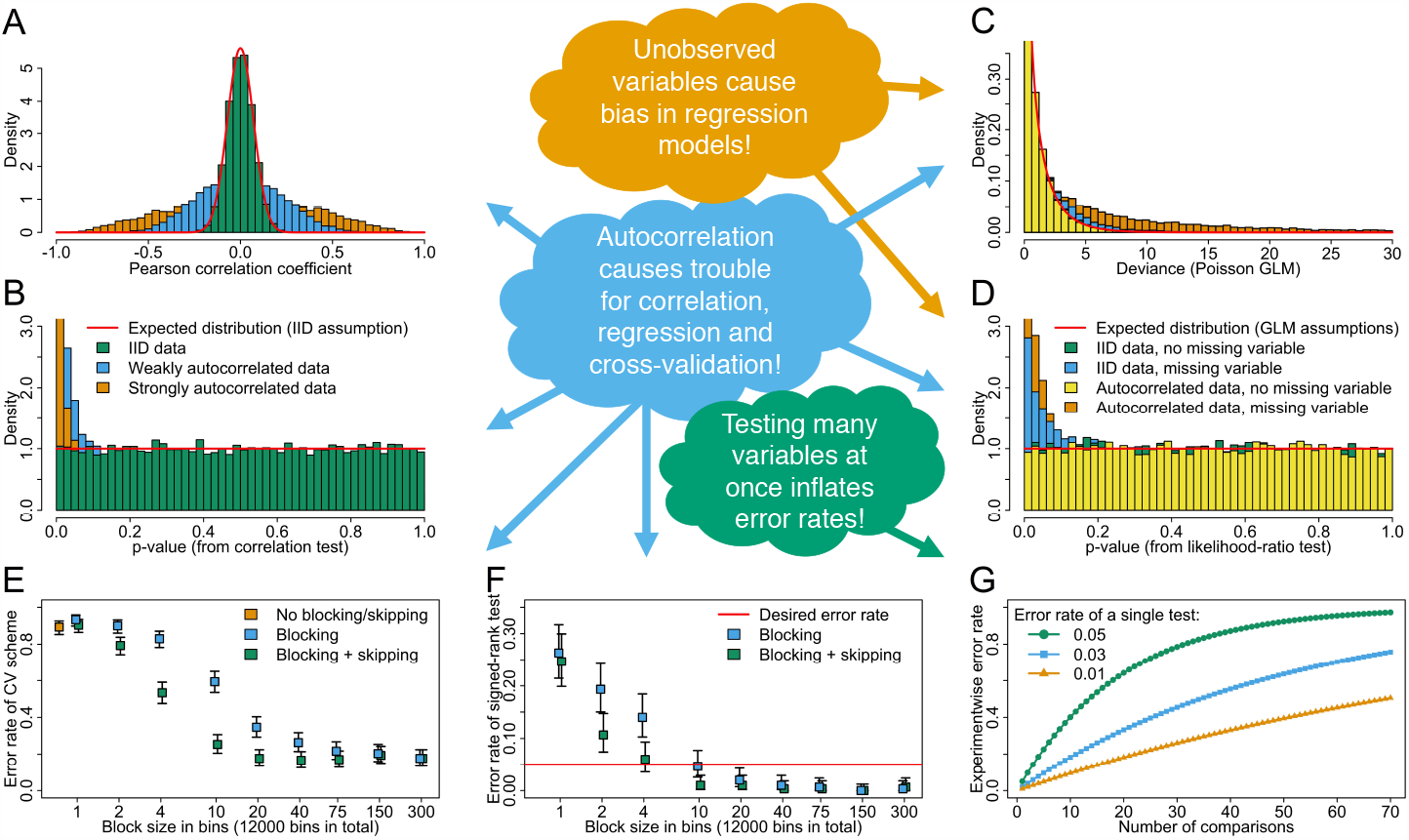
It is problematic to naively use statistics when necessary assumptions are not met. **A**: Empirical distribution of the Pearson correlation coefficient between two independent samples of 200 data points each, repeated 10000 times. Within sample the data are either independent or weakly or strongly autocorrelated. When the samples have temporal autocorrelation the empirical distribution of the correlation coefficient widens beyond the theoretical distribution for independent and identically distributed (IID) data. **B**: Corresponding distributions of *p*-values obtained under the assumption that the data is independent, showing that this mistake inflates the number of small *p*-values, yielding false positives (type I error) beyond the significance level set by the researcher. **C**: Empirical distribution of the deviance of a Poisson GLM comparing two nested models where one includes an irrelevant covariate (*x*-axis is cut at 30). 200 data points are generated of each covariate and the response variable, and this is repeated 10000 times. The covariates are either IID or autocorrelated, and we compare cases where there is or is not another relevant covariate missing from the model. **D**: Corresponding distributions of *p*-values obtained from likelihood-ratio tests comparing the nested models. When there exists a missing covariate, the number of small *p*-values and hence the rate of false positives are inflated. **E**: Type I error rates (how often does an overly complex model attain the best CV-score) for different CV schemes using different choices of block size and whether to skip blocks or not, including a scheme where folds are drawn completely at random, ignoring temporal structure. We see that failure to account properly for temporal dependencies greatly increases the probability that the wrong model is selected. Data is generated as in Results on simulated data, with 300 spike trains and 3 covariates simulated. The bars show 95% Clopper-Pearson confidence interval for the proportion of incorrect conclusions for each CV scheme. **F**: Same scenario as in **E**, except here a combination of cross-validation and the Wilcoxon signed-rank test is used. The significance level is set to *α* = 0.05, meaning that is the type I error rate we expect to see. With too small blocks the false positive rate is inflated, and with larger blocks the method appears overly conservative. **G**: Expected experimentwise type I error rates when performing multiple, independent tests, illustrating the need for correction. For a hypothesis test that on its own has type I error rate *α*, the experimentwise type I error rate grows as 1 − (1 − *α*)^*n*^.

Missing variables is another issue that is very likely to apply to any modeling of neural activity, since it is improbable that we are able to observe and correctly represent every underlying variable that is driving the neuron in question. Parameter estimates and *p*-values for the regression models are biased in the event of omitted variables (***Stevenson, 2018; Dunn and Battistin, 2017; Das and Fiete, 2020***). These omitted variables are typically also temporally autocorrelated, which further introduces temporal structure that could be misattributed to other variables. Figure 1 C and D show that missing variables substantially alter the distribution of the deviance of a GLM. It is illustrated specifically that a likelihood-ratio test based on GLM assumptions will behave as intended if all relevant variables are accounted for, but that the omission of relevant variables, autocorrelated or not, yields increased error rates.

The challenges mentioned so far rule out the use of the standard approaches in statistics for evaluating a given model performance, such as Akaike and Bayesian information criteria, which, similar to the likelihood-ratio test described above and illustrated in Figure 1 C and D, would yield overly optimistic results. Cross-validation (CV) is a commonly used method for model selection in such situations, estimating model performance by testing on withheld data folds (***Hastie et al., 2001***). To incorporate temporal structure, CV can be adapted with blocking (keeping blocks of consecutive time bins intact when constructing the folds) and skipping (removing portions of the data at the edges of the folds) (***Racine, 2000; Roberts et al., 2017***) (see Figure 9). If arbitrary data points or small blocks are used for constructing the folds, the error rate will be inflated (see Figure 1 E). The forward selection approach used by ***Hardcastle et al. (2017***) combines blocked CV with the Wilcoxon signed-rank test (***Wilcoxon, 1945***), a hypothesis test more general than those based on the GLM assumptions. Using small blocks, however, in the CV scheme may still inflate error rates in this approach (see Figure 1 F). Furthermore, testing only the best model in each step is a form of selective inference (***Taylor and Tibshirani, 2015***), akin to a multiple comparisons problem (***Goeman and Solari, 2014***), which can result in inflated false positives and requires correction (see Figure 1 G). If a small number of features are considered, however, the method tends to be conservative due to the general nature of the signed-rank test. This inflates the rate of false negatives, making it more challenging to discover actual relationships. Thus this approach can be overly permissive or very conservative depending on the data and the way the model is applied.

We evaluate different methods for covariate selection for the GLM, based on the forward selection procedure used by ***Hardcastle et al. (2017***), and explore possible adjustments to the CV scheme and hypothesis test used, in order to consolidate the inherent conservativeness of the hypothesis test and the lack of correction for multiple comparisons (note they do have a countermeasure for this, which we discuss briefly in Methods and Materials). We make modifications to their procedure by i) using skipping in the CV scheme to further separate the test and training data, ii) adjusting the number of folds, iii) considering different options for the hypothesis test used, and iv) correcting these tests for the multiple comparisons that are performed in each step of the forward selection procedure. The results we obtain with a modified Wilcoxon signed-rank test are similar to those of the unmodified one, while using a permutation test with cyclical shifts (details in Methods and Materials) yields higher test power than the signed-rank test, but with an increased computational cost. Finally, we discuss how the results of such studies, even when done as correctly as possible, can still be misleading, and suggest a way forward.

Our analyses are performed on simulated data and on calcium data from the medial entorhinal cortex (MEC) in freely moving mice (***Zong et al., 2022***). The methods are implemented in the statistical programming language R ***R Core Team*** (***2019***), and both simulated and real case studies were run on x86_64-apple-darwin15.6.0 (64-bit) with R version 3.6.0. All relevant code can be found at https://github.com/fredrine/covariateselection.

## Results

The methods that we have evaluated are summarized in Table 1. We denote the unmodified forward selection procedure using the Wilcoxon signed-rank test as SR and its Bonferroni corrected (for multiple testing) counterpart as SR_Bonf_. For these the *p*-values were calculated from a theoretical distribution of the test statistic. The procedure using modifications to the cross-validation (CV) scheme, as well as a maxT correction (***Westfall and Young, 1993***) for multiplicity is denoted as mSR_MaxT_. A further modified version of this where the new model were compared not to the previous one, but one also including the added covariate reversed in time, is denoted as mSRR_MaxT_. To able to use the maxT correction we calculated the *p*-values by constructing a shuffled distribution by permuting the signs of the CV scores for these methods. CS_Bonf_ is used to denote the procedure using the modified CV scheme, with a permutation test with cyclical shifts in place of the signed-rank test. Cross-validation, with blocking and skipping, used to directly select covariates without any hypothesis test is denoted as CV. Detailed descriptions of all the methods can be found in Methods and Materials. Figure 2 illustrates how a distribution of *p*-values will look when the performed test is either invalid, valid or conservative.

**Table 1.** Summary of different hypothesis tests used in step 3 of the forward selection procedure, and the corresponding CV scheme used to determine which covariate that should be added to the model.

| Method | Test statistic | Distribution | Correction | Skipping |
| --- | --- | --- | --- | --- |
| SR | Signed-rank new vs. previous | Theoretical | None | No |
| SR<sub>Bonf</sub> | Signed-rank new vs. previous | Theoretical | Bonferroni | No |
| mSR<sub>MaxT</sub> | Signed-rank new vs. previous | Perm. of signs | MaxT | Yes |
| mSRR<sub>MaxT</sub> | Signed-rank new vs. reversed | Perm. of signs | MaxT | Yes |
| CS<sub>Bonf</sub> | Log-likelihood new vs. previous | Cyclic shift perm. | Bonferroni | Yes |

**Figure 2.**
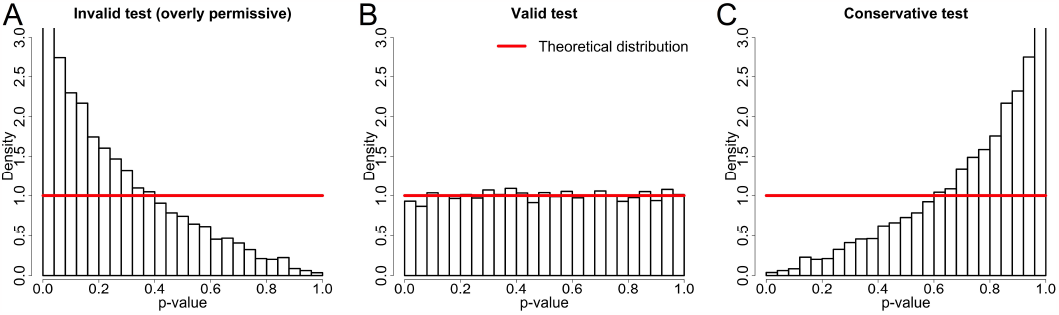
Different hypothetical distributions of *p*-values resulting from a hypothesis test. For the test to be valid, the probability of obtaining a *p*-value smaller than a specified significance level *α* must be at most *α*, i.e. Pr(*p* ≤ *α*) ≤ *α*, when the null hypothesis is assumed to be true. If this is not true, and Pr(*p* ≤ *α*) > *α*, the distribution will be skewed towards 0, the rate of false positives will be inflated (**A**), and the test is invalid. For a valid test we typically see Pr(*p* ≤ *α*) = *α* (**B**), giving a uniform distribution of *p*-values. A test might also be conservative, meaning it is still valid, but the distribution is skewed towards 1 (**C**). In the latter case the rate of false negatives will be inflated, and it is less likely that true positive results are discovered.

### Results on simulated data

We have conducted a simulation study to test the different methods under conditions similar to real data, but where we know the underlying truth. Variables with temporal autocorrelation are constructed by passing a vector of independently sampled uniform variables through a smoothing filter. One such variable is treated as unobserved, but driving the cell to fire. Three other variables are sampled independently from the driving variable and each other, and are treated as potential covariates for the regression model. One of these is generated as a two-dimensional variable, emulating animal position.

Two different scenarios are simulated. In scenario 1, the unobserved variable is all that drives the simulated cell, and thus the observed variables are all irrelevant. In this case we would like our methods to conclude that no variable should be included in the model. Furthermore, for a hypothesis test to be valid the probability of observing a *p*-value below the significance level *α* (for any choice of 0 < *α* < 1) must not exceed *α* (***Casella and Berger, 2002***, p.397). In practice this means that the distribution of observed *p*-values should be uniform or skewed to the right. The latter could however be an indication that the hypothesis test is conservative, and potentially lacking in power. In scenario 2, the cell activity is also affected by the two-dimensional variable, specifically by having heightened spike activity in two specific spatial positions. In this case our methods would ideally recognize that the two-dimensional variable should be included in the model, and that the others should not.

Figure 3 shows the results from the forward selection procedure in scenario 1, where none of the observed covariates are relevant to the cell activity. In part A there are histograms showing how often each of the possible covariate combinations is selected as the final model for each of the methods evaluated. The methods all conclude that few to no covariates should be included. CV alone (no hypothesis test conducted) naturally yields more complex models than the other methods, and is the only method with a type I error rate exceeding the selected significance level of *α* = 0.05. The remaining methods all have reasonable error rates, where mSR_MaxT_ and SR_Bonf_ in particular appear overly conservative with no false positives. The modified signed-rank test testing against the model with the reversed covariate (mSRR_MaxT_) and the unmodified signed-rank test (SR) both appear conservative, while the method using cyclic shifts (CS_Bonf_) has an error rate closer to the selected significance level. A summary of the error rates with corresponding 95% Clopper-Pearson confidence intervals are shown in Table 2.

**Table 2.** Summary of results when using simulated data with 300 simulated cells. We show type I error rates (proportion of simulations where a covariate is incorrectly included) for scenario 1, where none of the observed covariates affect the response, and power (proportion of simulations where the two-dimensional covariate is correctly included) for scenario 2, where the two-dimensional covariate has an effect on the response. We also show corresponding 95% Clopper-Pearson confidence intervals. For each hypothesis test the significance level *α* = 0.05 is used.

| | No effect | $n = 300$ | 2D effect | $n = 300$ |
| --- | --- | --- | --- | --- |
| Method | Error rate | 95% CI | Power | 95% CI |
| CV | 0.220 | (0.174, 0.271) | 1 | (0.988, 1.000) |
| mSR<sub>MaxT</sub> | 0 | (0.000, 0.012) | 0.443 | (0.386, 0.502) |
| mSRR<sub>MaxT</sub> | 0.010 | (0.002, 0.029) | 0.690 | (0.634, 0.742) |
| CS<sub>Bonf</sub> | 0.037 | (0.018, 0.065) | 0.993 | (0.976, 0.999) |
| SR | 0.003 | (0.000, 0.018) | 0.627 | (0.569, 0.682) |
| SR<sub>Bonf</sub> | 0 | (0.000, 0.012) | 0.297 | (0.246, 0.352) |

**Figure 3.**
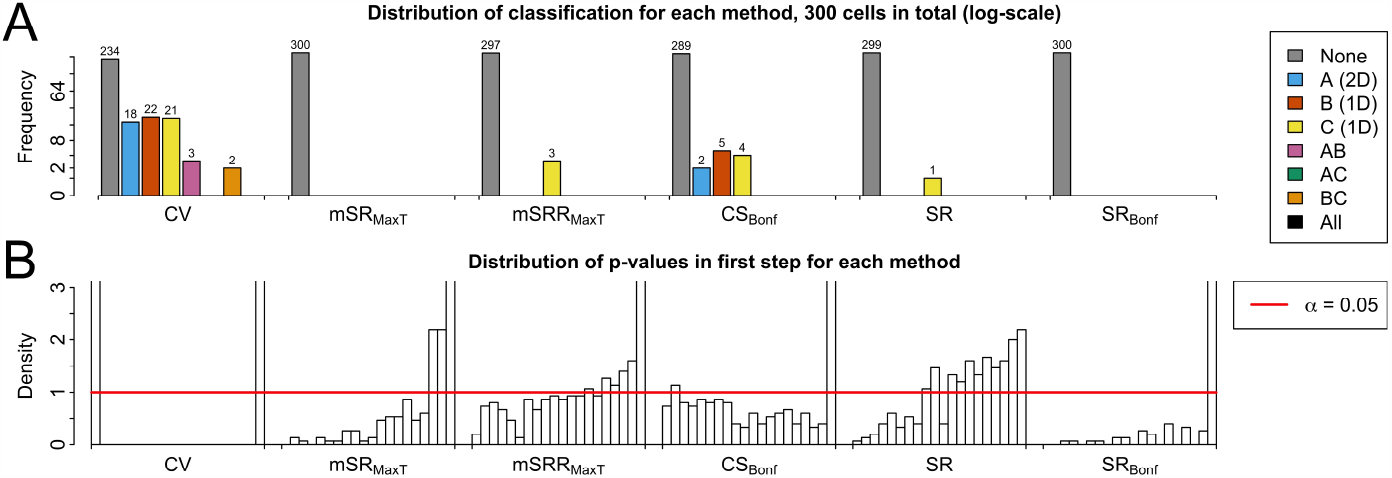
Testing simulated data where none of the observed covariates have any effect on the response. **A**: For each method (CV, mSR_MaxT_, mSRR_MaxT_, CS_Bonf_, SR, SR_Bonf_, see Table 1) the number of cells classified as tuned to each of the 8 combinations of the three covariates is shown, for a total of 300 simulated cells. The bars are colored according to which final model is selected by the forward selection procedure. Note that the y-axis is in log-scale to emphasize sparse results. **B**: Distribution of *p*-values for each method at the first step. For this scenario the proportion of *p*-values smaller than *α* = 0.05 corresponds to the type I error rate, and must itself be smaller than *α* on average for the method to be valid. CV without any added hypothesis test exceeds this threshold, whereas the other methods all give error rates below it.

The results from scenario 2, where the simulated two-dimensional covariate affects the cell activity, is shown in Figure 4, and summarized in Table 2. The correct covariate is identified by the CV scheme in every simulation. The method using the cyclical permutation test (CS_Bonf_) has the highest estimated test power (defined here as the proportion of simulations where the two-dimensional covariate is correctly included) of the methods which include a hypothesis test. We see smaller test power for mSR_MaxT_ than its alternative counterpart mSRR_MaxT_. We can observe that the results from SR are similar to mSRR_MaxT_, suggesting that the increased power from comparing to the model with the reversed covariate is on par with the increased power from not accounting for multiple comparisons, in this case. Generally the latter will likely scale differently with the number of covariates considered, which in the extreme case will cause inflated type I errors (see Figure 1). For SR_Bonf_ we observe low power, in line with how it appears conservative in Figure 3.

**Figure 4.**
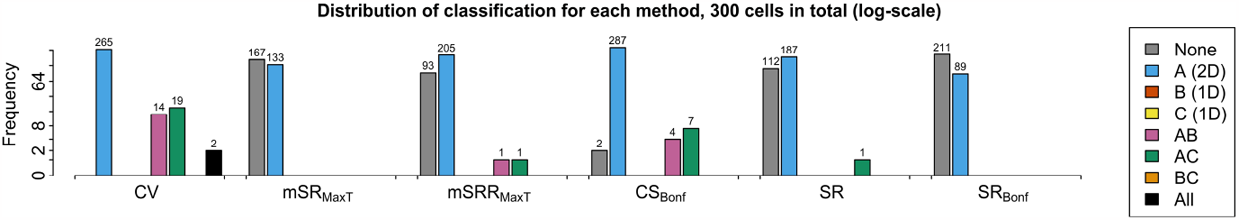
Testing on simulated data where the simulated position covariate has an effect on the response. For each method (CV, mSR_MaxT_, mSRR_MaxT_, CS_Bonf_, SR, SR_Bonf_, see Table 1) the number of cells classified as tuned to each of the 8 combinations of the three covariates is shown, for a total of 300 simulated cells. The bars are colored according to which final model is selected by the forward selection procedure. We see that the methods in general correctly favor the pure position model. Note that the y-axis is in log-scale to emphasize sparse results. Covariate A is relevant to the activity of the simulated cells and should be included in the model. Additionally, since we use a significance level of *α* = 0.05, we expect to see additional, irrelevant covariates for approximately 5% of the cells where the correct one is included.

### Results on calcium data

A study using calcium data from the medial entorhinal cortex (MEC) of freely moving mice is also conducted to test the methods on real data. The data is the publicly available dataset from ***Zong et al. (2022***) and ***Obenhaus et al. (2022***), who classify grid cells using the cyclical shift permutation test by shifting the neural activity in time, and the grid score as the test statistic.

We test the methods again in two scenarios. In scenario 1, we combine cell activity from one session (approximately 27 minutes) with the behavioral covariates of another, so that the observed variables are all irrelevant to the response. Thus we expect the methods to conclude that no variable should be included in the model. In scenario 2, the cell activity is matched with the correct behavioral covariates, and thus we expect to see the methods classify the cells as having tuning. We do not however know the ground truth in this scenario, but we can assess how well our methods identify the grid cells previously classified by ***Zong et al. (2022***) as position-tuned.

The results from scenario 1, with mismatched cell activity and covariates, are shown in Figure 5, tested with 249 cells. A summary of the error rates can be found in Table 3. All methods select few to no covariates, similarly to what we observed in scenario 1 with simulated data. CV alone yields the most complex models, and thus has the highest error rate. The unmodified signed-rank test (SR), the Bonferroni corrected version of this (SR_Bonf_), and the maxT corrected version where we perform the permutation of signs (mSR_MaxT_) all appear overly conservative, yielding no false positives. The alternative version of the latter (mSRR_MaxT_) and the cyclic shift based method (CS_Bonf_) commit slightly more errors, but not more than the selected significance level of *α* = 0.05.

**Table 3.**
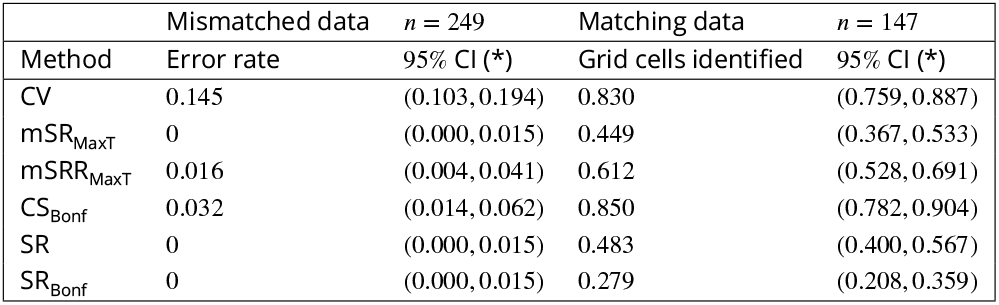
Summary of type I error rates (proportion of cells where a covariate is incorrectly included) for the scenario with mismatched data where none of the observed covariates affect the response, and proportion of the 147 grid cells classified by ***Zong et al. (2022***) that are also classified as position tuned. We also show corresponding 95% Clopper-Pearson confidence intervals. For each hypothesis test the significance level *α* = 0.05 is used. (*) The confidence intervals are based on the binomial distribution, which assumes that the neurons are independent. In reality there are small correlations present, meaning that the intervals are slightly too optimistic, and should be somewhat wider.

**Figure 5.**
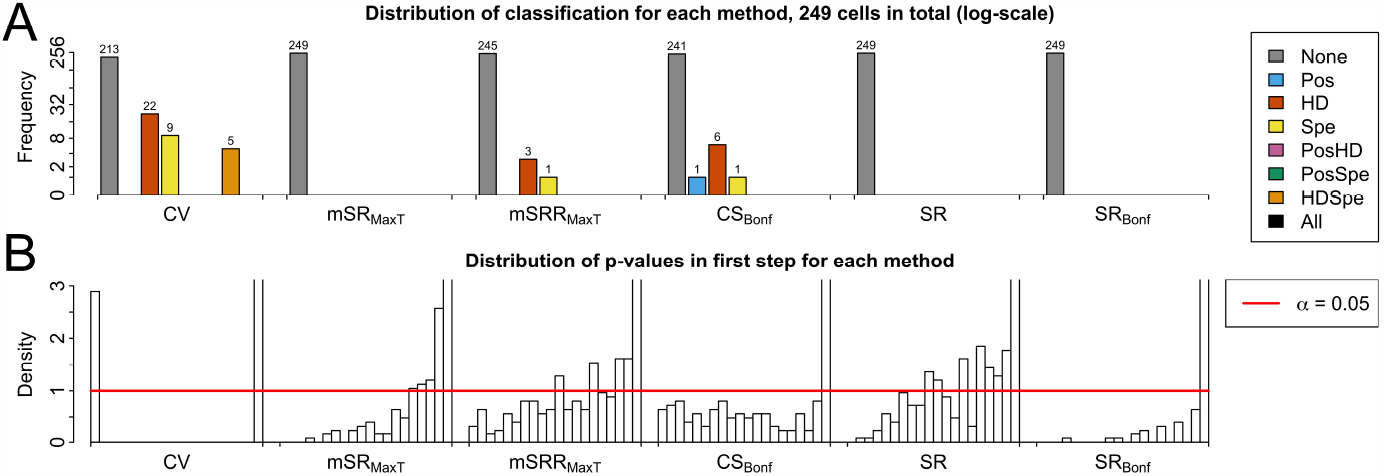
Results from testing on calcium data where the cell activity and covariates are from different sessions. **A**: For each method (CV, mSR_MaxT_, mSRR_MaxT_, CS_Bonf_, SR, SR_Bonf_, see Table 1) the number of cells classified as tuned to each of the 8 combinations of the three covariates is shown, for a total of 249 cells. The bars are colored according to which final model is selected by the forward selection procedure. Note that the y-axis is in log-scale to emphasize sparse results. **B**: Distribution of *p*-values for each method at the first step. For this scenario the proportion of *p*-values smaller than *α* = 0.05 corresponds to the type I error rate, and must itself be smaller than *α* on average for the method to be valid. Cross-validation without any added hypothesis test exceeds this threshold, whereas the other methods all give error rates below it.

The confidence intervals included in Table 3 are constructed in the same way as in Table 2 in the previous section, using Clopper-Pearson method for a binomial proportion. However, while the simulated neurons in the previous section are generated independently, the activity of actual neurons will be correlated to some extent. Therefore the assumption of independence between the neurons, which is made when using the binomial distribution, is violated. The confidence intervals in Table 3 are therefore likely to be overly optimistic, meaning they are too narrow. Methods that account for dependencies are likely to be overly conservative since the correlations are small and limited to clusters of neurons. A solution to this issue which accounts more specifically for the type of correlation structure that is found in neurons is to our knowledge not found in the field.

Figure 6 shows the results from our tests on the calcium data in scenario 2, where the cell activity and covariates are correctly matched. Most cells are classified as position-tuned by all the methods, while CV and CS_Bonf_ both suggest that head direction or speed also might have an effect on the cell activity of a large portion of the cells. Table 3 shows for each method evaluated the proportion of the 147 grid cells classified by ***Zong et al. (2022***) that the GLM also classified as position tuned. CV and CS_Bonf_ classify the most grid cells, followed by mSRR_MaxT_, while the remaining methods classify less than half of these cells.

**Figure 6.**
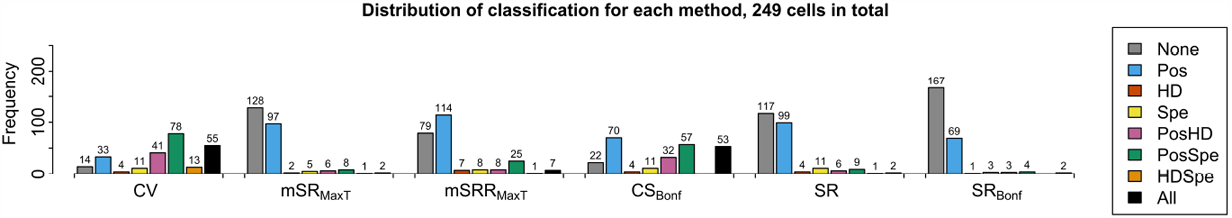
Results from testing on calcium data where the cell activity and covariates are from the same session. For each method (CV, mSR_MaxT_, mSRR_MaxT_, CS_Bonf_, SR, SR_Bonf_, see Table 1) the number of cells classified as tuned to each of the 8 combinations of the three covariates is shown, for a total of 249 cells. The bars are colored according to which final model is selected by the forward selection procedure. Most methods favor models including position as a covariate, and CS often suggests a model with multiple covariates.

Ratemaps for some of the grid cells that CS_Bonf_ classifies as position tuned are shown to the left in Figure 7. The middle of the same figure shows ratemaps of some of the 87 cells classified as position tuned by the GLM, but that are not classified as grid cells. For most of these cells the ratemaps convincingly indicate that there is a structural relationship between firing and position, and some even show weak grid patterns, suggesting they barely failed to be classified as grid cells. The rightmost part of the figure shows ratemaps for 9 of the 22 cells that are classified as grid cells by ***Zong et al. (2022***), but not recognized as position tuned by the GLM. We can see that the firing patterns of these cells generally appear more foggy than those in the leftmost plot, and the average mean event rate for these neurons are also smaller than the ones that are classified by the GLM, which might explain why these are not. The grid score is also a more specialized way to classify neurons than the GLM, meaning weaker patterns may still be found by the former because it specifically looks for those patterns.

**Figure 7.**
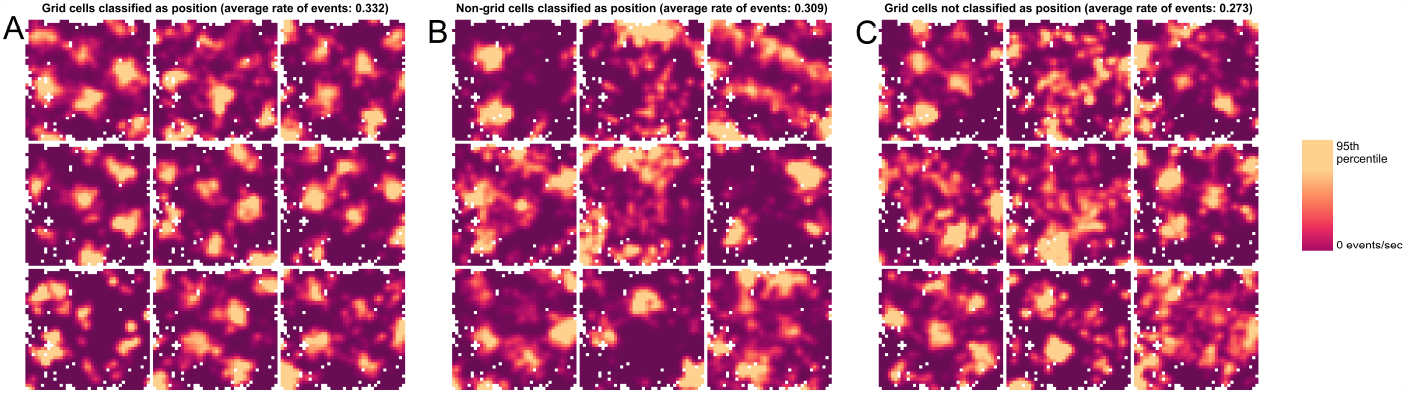
Ratemaps and tuning curves for a selection of cells with different classifications by the GLM approach with CS_Bonf_. **A**: Cells classified as grid cells by ***Zong et al. (2022***), and as position tuned by the GLM. **B**: Cells not classified as grid cells, but as position tuned by the GLM. **C**: Cells classified as grid cells, but not classified as position tuned by the GLM. Animal position is binned using 40 bins in each direction, and the average number of events occurring in each two-dimensional bin is calculated. Then a two-dimensional Gaussian filter with standard deviation of 1 bin is applied. Each ratemap is then normalized by dividing it by its 95th percentile. The mean of the average event rates for each group the subset of cells is from is displayed in each plot title.

Figure 8 shows the number of cells that mSRR_MaxT_, SR and CS_Bonf_ classify as tuned to the different covariates, as well as the number of grid cells classified by ***Zong et al. (2022***). Most of the 147 grid cells are classified with each method, with CS_Bonf_ classifying the most. As with the simulated data we can also here observe that the results from SR are similar to those from mSRR_MaxT_. When using CS_Bonf_ we obtain results suggesting the cells have conjunctive tuning to two or even all three of the covariates considered, with 53 cells (21.6%) being classified as tuned to all three. We do not know the underlying truth of what is driving the firing of the neurons, but these results are consistent with other findings in the MEC (***Sargolini et al., 2006; Hardcastle et al., 2017***).

**Figure 8.**
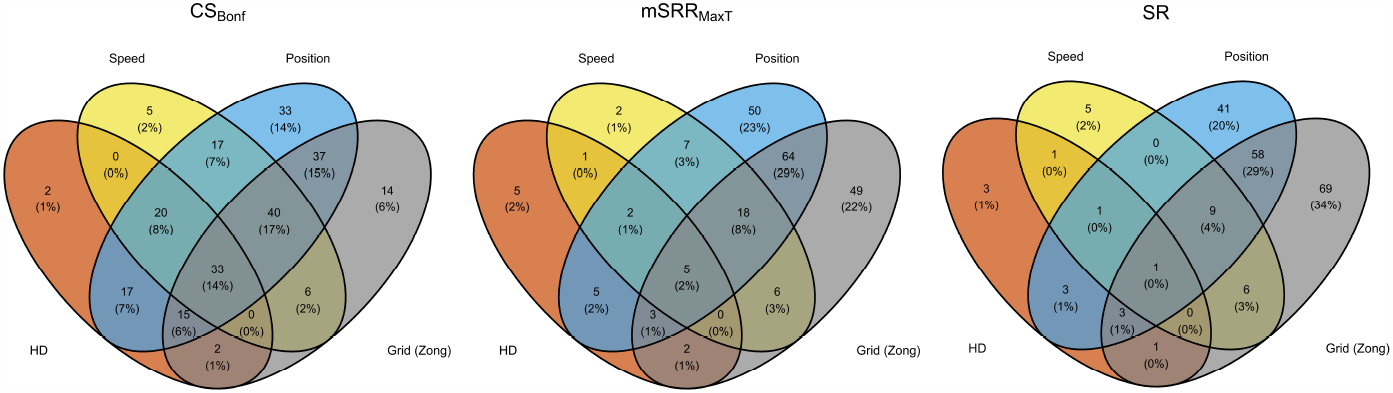
Resulting classification from the GLM using CS_Bonf_, mSRR_MaxT_ and SR, compared to the grid cell classification by ***Zong et al. (2022***). This means that CS_Bonf_ classifies for example 2 + 0 + 20 + 17 + 33 + 15 + 2 + 0 = 89 cells as tuned to head direction, 17 + 20 + 16 + 31 + 37 + 40 + 33 + 15 = 209 to position, 33 + 20 = 53 as tuned to position, head direction and speed, and that 37 + 40 + 33 + 15 = 125 of the 147 grid cells are classified as position-tuned. In general CS_Bonf_ classifies more cells than the other two methods, which yield similar results.

**Figure 9.**
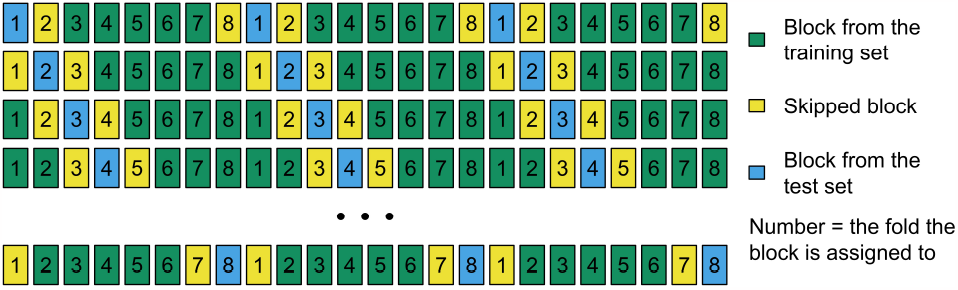
Illustration of CV scheme with blocking and skipping. Each box is a block of consecutive time bins. When a fold is used as the test set, the neighboring blocks are excluded from the training set. Note that there are 8 folds in the illustration, compared to 10 or 20 in our analyses.

## Discussion

We show through testing on both simulated data and real neural data that methods typically used for covariate selection in neuroscience can be either overly conservative or overly permissive, but they can be made statistically robust and even more powerful with a few modifications.

Cross-validation (CV) on its own can work well for the purpose of covariate selection if the temporal structure of the data is taken into consideration, although it typically yields false positive rates beyond desirable levels. This might be sufficient if the goal of the analysis is to explore, rather than decisively conclude which covariates are relevant to the neural response. If the analysis is performed with the intent of conducting confirmatory research, additional hypothesis testing in the forward selection scheme is desired to control the rate of false positives.

The Wilcoxon signed-rank test is inherently conservative when testing CV log-likelihood values of nested models, meaning that their type I error rate control is stricter than the threshold set by the user (explained further in Methods and Materials). If the method is used incorrectly, however, either by using too small blocks of data for the CV, or by failing to account for multiple testing, the false positive rate may instead be inflated (see Figure 1). The modification of the signed-rank test where we compare the proposed model to one where the added covariate is reversed alleviates some of the conservative tendencies of the test, because we compare models of equal complexity. We see that this results in increased test power both for simulated and real data.

The forward selection procedure without modifications to the CV scheme and signed-rank test (SR) yields very similar results to our adjusted method (mSRR_MaxT_), so for this particular case it may seem like the inherent conservativeness and lack of correction for multiple comparisons cancel each other out, coincidentally. We should however strive for statistical robustness and generality, meaning the adjusted method should be preferred, as statistics of each data set will be different. The permutation test using cyclical shifts of the covariate considered for the model attains desirable error rates and higher test power than the other methods. This does however come at computational cost.

An additional remark with respect to statistical robustness is that even though the methods using the signed-rank test appear valid, we should be cautious. The fact that the CV training sets overlap to some extent, potentially violating signed-rank test assumptions, means that we are unable to completely trust the generality of the results. This issue was addressed by ***Dietterich*** (***1998***), who proposed a solution of using a 5 x 2 CV test, where the data is split into CV folds multiple times in order to reduce the effect of overlapping training sets. However, with temporal data, such as ours, the repeated splitting of the data is not feasible unless the data length is very long.

The methods presented in this work serve as a proposal to the problems caused by temporal structure without having to model this structure specifically. Including past spike history using auto-regressive terms in the regression model is common in the field (***Truccolo et al., 2005***). While these terms have biophysical interpretations at smaller time scales, where they for example can represent the refractory period of the neuron, they are unlikely to remove all temporal dependency due to unobserved, temporally dependent factors also driving the activity (***Stevenson, 2018***). Hence the additional adjustments we suggest are warranted.

An overarching goal in neuroscience is to determine which purpose the brain area in question serves, and classifying individual neurons from the area in terms of tuning to covariates is one possible approach. If we treat the classification of each neuron as a vote, we can likely conclude that the covariates with the most votes are important to the brain region as a whole. Furthermore, an anatomical clustering of cells with similar tuning is logical in the sense that these cells rely on input from each other to function as a network. ***Obenhaus et al. (2022***) argue that this is true in particular for the medial entorhinal cortex (MEC), where a mental map is constructed by different cell types all related to navigation.

Not all cells are specialized into cell types, however, and instead of modelling single neurons another possibility is to use methods and models that consider the population code more directly, using for example latent variable models (LVM) (***Lawrence, 2003; Yu et al., 2008; Wu et al., 2017; Pandarinath et al., 2018; Bjerke et al., 2023; Schneider et al., 2023***) or topological data analysis (TDA) (***Curto and Itskov, 2008; Rybakken et al., 2019; Gardner et al., 2022***). Such methods can be used without knowledge of specific covariates. However, single-cell classification is still useful, as it can be used to group neurons into coherent populations with a neural code which is likely correlated to the covariate observed (such as in ***Gardner et al. (2022***)).

Finally, we emphasize that it is important that the statistical methods used are robust, so that the results they yield can be trusted and replicated.

### Moving forward

Perhaps the most important aspect of these analyses is the data itself. When designing future experiments we would suggest both minimizing the number of possible candidates for covariates while also making these as complete as possible and recording for a long time. This occurs largely at the experimental design phase where care should be taken to limit the number of likely behaviors and engagement of different sensory-motor modalities. For example, simply turning off the lights removes a complex and difficult-to-track set of covariates, while engaging the animal actively in a specific task such as pursuit reduces the number of spurious behaviors that might not occur often enough to be properly studied. Second, it is worth the effort to properly track and invest in constructing compact postural and behavioral variables that account for the strong correlations between, for example, how the head and the body rotate together (***Mimica et al., 2018***). Third, the experiment should ensure that each covariate is sampled well. Untangling the potential spurious relationships between our covariates is important to improve our ability to interpret the results we obtain. In this regard we should strive for long experiments with rich and diverse behavior, to ensure the covariate space is properly explored. Longer data sets will also make it easier to account for temporal autocorrelation, as it allows for larger blocks of data for methods such as cross-validation.

With a minimal but hopefully close to a complete set of covariates, we would suggest using the cyclical shift permutation with the general Bonferroni correction for covariate selection. The other methods we study here are computationally more efficient, however, they can either be much more permissive or conservative than expected. If there is reason to be believe that the data is not stationary, an alternative to the cyclical shift should be used. If one suspects that there is a trend throughout the experiment in either behavior or neural activity, the linear shift test proposed by (***Harris, 2020b***) will be more appropriate. An option for trial-based experiments is the session permutation method described by ***Harris*** (***2020a***), where the shuffled distribution is constructed by repeatedly calculating the test statistic with neural data from the trial considered and behavioral variables from a trial selected randomly from the remaining ones.

If the computational cost of the cyclical shift is too high, we recommend the signed-rank test with the modifications to the cross-validation scheme and testing the new model against the one with the reversed covariate (mSRR_MaxT_). However, the extent to which the overlapping data affects the false positive rates should be further explored, and suitable modifications to account for potentially inflated error rates should be found.

We also suggest checking the false positive rates, regardless of the approach or model, by swapping the covariates from different experiments and constructing simulations. This does not ensure that the model is correct but is a simple way of showing that a seemingly significant result is indeed not (e.g. the case of potential mirror-like cells in mice (***Tombaz et al., 2020***)).

We stress again that interpretation of results is difficult, as these can be misleading in situations where there are strong task-dependent relationships between different covariates that may change depending on the behavior of the animal. Unobserved, but relevant variables further complicate the matter. A cell appearing tuned to speed might for instance very well be driven by a specific behavior that only occurs when the animal is moving at certain speeds. Thus when one observes a prominence of tuning to a selection of covariates, it is important to consider what alternative and possibly more likely explanations might exist. This issue is not unique to the GLM framework, and is something that should be taken into consideration when we draw our conclusions regardless of approach.

## Methods and Materials

### Model framework

We consider recordings of neural activity and tracked external covariates over an interval of time. When using a generalized linear model (GLM) the response variable *Y*_*i*_, *i* = 1, 2, …, *n*, are assumed to be random variables from a distribution in the exponential family, typically either the number of spikes in a time bin, using the Poisson distribution, or a binary variable representing cell being inactive or active, using the Bernoulli distribution. The expected value of *Y*_*i*_ conditioned on observing the vector of covariates *X*_*i*_ = *x*_*i*_ is specified using a link function and functions of the covariates. More specifically,

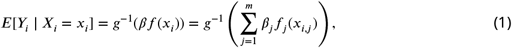

where *g* is the link function connecting the expected value *μ* and the linear predictor *η* and *f* is an optional function of the covariates. We use a Bernoulli GLM with logit-link, 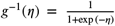 Through the functions *f*_*j*_ the covariates can be represented in various ways, for example using binning. We represent our covariates by cubic splines.

In the GLM the responses are additionally assumed to be independent when conditioned on the covariates included in the model, which is unlikely to hold for temporal neural data. The temporal autocorrelation in the *Y*_*i*_ might in part be explained by biophysical properties of the neuron at high temporal resolution (time bins with widths of a few milliseconds) (***Truccolo et al., 2005***), but will also to a large extent be caused by the omission of relevant covariates (***Stevenson, 2018***). See Figure 1 for a demonstration of how this model misspecification, and generally naive use of statistics, can cause trouble.

### Forward selection procedure

We use a forward selection procedure similar to that of ***Hardcastle et al. (2017***), which is based on cross-validated (CV) log-likelihood values and performing a hypothesis test at each step. Starting with a model without covariates (hence modelling a constant mean firing rate), the procedure goes as follows

1. Perform CV to find the covariate that improves the model the most by comparing the mean difference of paired CV log-likelihood values
2. Perform a hypothesis test, where the null hypothesis is that the added covariate does not improve the model (i.e. the expected value of the mean difference of paired CV log-likelihood values is 0)
3. If the new model is significantly better than the previous one, continue from step 1, otherwise the previous model is chosen as the final one

### Cross-validation scheme

Cross-validation is a popular method for model selection in many fields, which seeks to estimate model performance on held out data (***Hastie et al., 2001***). The data is partitioned into folds, where each fold once serves as a test set while the model is trained on the remaining folds. Caution is required for data with temporal structure, as otherwise each fold might contain highly correlated data points, resulting in an overly optimistic test performance.

One common approach to account for this is to temporally block the data points and construct folds of whole blocks (***Roberts et al., 2017***). In addition one might want to isolate the test fold further, meaning that we remove a few of the neighboring data points from the training set (***Racine, 2000***), illustrated in Figure 9.

In our main analyses on simulated and real data we use *N* = 12000 data points. In the modified CV scheme we use 20 folds, where each fold consists of 4 temporally separated blocks, each thus of 150 consecutive time bins. These 20 ⋅ 4 = 80 blocks are systematically chosen to belong to the 20 folds, meaning the first, 21st, and 41st block belong to fold 1, and so on. For the unmodified CV scheme we use 10 folds and blocks of 150 time bins, meaning each fold consists of 8 blocks. We also use skipping, and exclude folds *i* + 1 and *i* − 1 from the training set when using fold *i* as a test set, meaning the 150 time bins on each side of the blocks belonging to fold *i* are excluded. When fold 1 is used as a test set, fold 20 is excluded from the training set (and vice versa). The last (first) block is also then excluded even though it does not border any of the blocks from fold 1 (20), in order to keep the length of the training sets consistent.

***Hardcastle et al. (2017***) use 10 folds, which we thus also use for the methods with the un-modified CV scheme (SR and SR_Bonf_). For the methods with a modified CV scheme (mSR_MaxT_ and mSRR_MaxT_) we use 20 for two reasons. Firstly, with the added step of skipping neighboring blocks for the CV, more folds means more data is available in each training set, yielding a less biased estimate of the mean difference in log-likelihood values. Secondly, the signed-rank test statistic, and thus the corresponding *p*-value, can only take a finite number of values (2^*k*^ with *k* folds), which with *k* = 10 scales poorly when you correct for multiple comparisons.

### Permutation test procedure

Permutation testing can be used when performing a hypothesis test where the distribution of the test statistic under the null hypothesis is not known (***Westfall and Young, 1993***). The key property that makes this possible is exchangeability, meaning that the distribution of a vector of random variables is unchanged by a permutation of the indices. This property is not fulfilled for arbitrary permutations when the random vector has temporal structure, since this is distorted. Instead it is common in the field (***Grijseels et al., 2021; Zong et al., 2022; Høydal et al., 2019***) and also fields involving spatial dependencies (***Dale and Fortin, 2002***) to use a restricted subset of possible permutations consisting of cyclical (toroidal in the spatial case) shifts for randomly drawn lags, illustrated in Figure 10.

**Figure 10.**
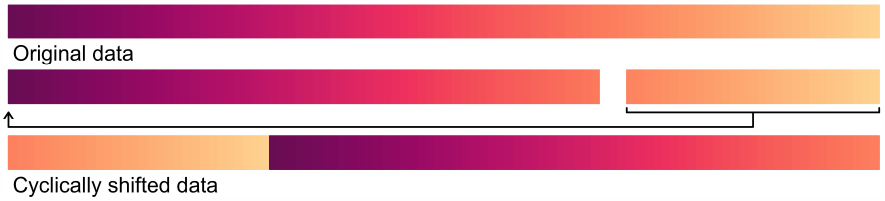
Illustration of the cyclical shift permutation.

The procedure is thus as follows for some permutation test:

1. Calculate the test statistic *t*
2. Perform *B* permutations to generate a null distribution 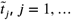, *B*
3. Calculate permutation *p*-value as 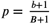, where 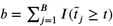 is the number of times the permuted sample yields a test statistic larger than or equal to the originally observed one
4. If doing Bonferroni correction, calculate adjusted *p*-value as *p*_adj_ = *p* ⋅ *m*

Note the adjustment 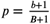 as opposed to 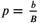, which we do to prevent nonsensically reporting a *p*-value of 0, and also because the latter is not valid (***Phipson and Smyth, 2010***).

### Hypothesis test based on the Wilcoxon signed-rank test

The signed-rank test methods used are based on differences of cross-validated log-likelihood values. We use *k* = 20 or *k* = 10 fold CV, yielding values *A*_*i*_, *i* = 1, …, *k* for the current model and values *B*_*i*_, *i* = 1, …, *k* for the candidate model. We wish to test if the *B*_*i*_ are significantly larger than the *A*_*i*_.

We consider the differences *D*_*i*_ = *B*_*i*_ − *A*_*i*_. The null hypothesis is then *H*_0_: the distribution of the *D*_*i*_ is symmetric about *μ* = 0, where *μ* is the expected value of each *D*_*i*_. The test statistic used is the signed-rank:

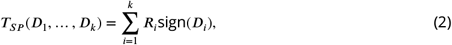

where *R*_*i*_ is the rank of the difference *D*_*i*_ when they are ordered by their absolute values, so that the smallest |*D*_*i*_| has rank 1 and the largest has rank *k*.

The observed test statistic is then either compared to a theoretical distribution (SR and SR_Bonf_), utilizing the `wilcox.test` function from the `stats` package in R, or to an empirical distribution constructed by permutations where we flip signs (mSR_MaxT_ and mSRR_MaxT_), meaning we use the permutation *ρ*(*D*_*i*_) = *u* ⋅ *D*_*i*_, where *u* is a random vector where each element is independently sampled from {−1, 1} with equal probabilities. We run 999 permutations for this method.

Note that under the null hypothesis the Wilcoxon signed-rank test assumes that the distribution of the CV log-likelihood differences are centered at 0. When comparing a model to the one with an additional and irrelevant covariate, however, the new model will overfit and thus generally yield negative CV log-likelihood differences. This means that the resulting test will be overly conservative (provided the CV is done properly, since improper CV can yield large values even when overfitting, see Figure 1.

Because of this we also consider the variation where the proposed new model is compared not to the previous one, but to a model where the additional covariate is included in reversed order (mSRR_MaxT_). Under the null hypothesis that the added covariate has no effect on the response, and assuming the distribution of the covariate to be stationary, these two models should explain the response equally well, and hence the differences in CV log-likelihood values will be centered around 0.

Improper CV will also violate the assumption of the signed-rank test that the differences themselves must be independent. Without skipping and without or with too small blocks, neighboring folds in the CV will be highly correlated, which also means that the CV log-likelihood differences will be, further motivating the need for these adjustments. Even with these adjustments, however, there is overlap between the training sets in each iteration of the CV, potentially inducing correlation between the CV log-likelihood differences. The signed-rank test does appear robust to this problem in both the simulated and real data analyses in this work, but we do not know under what conditions it will be a problem.

### Hypothesis test based on cyclical shifts

For the cyclical shift permutation test the in-sample log-likelihood increase is used as a test statistic, and the covariate that is being added is shifted cyclically to generate a distribution of this test statistic under the null hypothesis. The covariate vector (or each column representing it if the covariate has multiple parameters) is shifted in time by a random lag, and the part that is shifted beyond the original end of the session is wrapped to the beginning, illustrated in Figure 10. We thus assume that the distribution of the covariate added is stationary.

When comparing a current model (A) and model (B) with the added covariate, our test statistic becomes

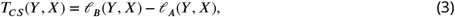

where *ℓ*(*y, x*) is the log-likelihood of the GLM with coefficients *β* for the Bernoulli case, i.e.

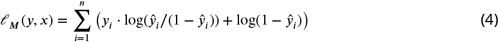

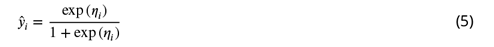

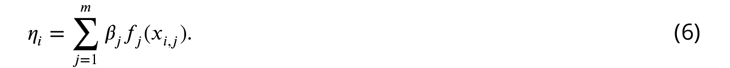

Without temporal structure and unobserved covariates, 2*T*_*CS*_(*Y, X*) (deviance of the Bernoulli GLM) asymptotically follows a *χ*^2^ distribution with degrees of freedom equal to the difference in number of parameters between the two models (***Wilks, 1938***). However, as we illustrated in Figure 1, this is not the case when these challenges are present, which motivates the use of a permutation test approach to estimate the actual distribution of the test statistic.

We permute by drawing a random lag *l* uniformly from {150, 151, …, *n* − 150}, so the permuted version of the covariate *x* = (*x*_1_, *x*_2_, …, *x*_*n*_) becomes 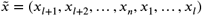. The permutations are more costly with the cyclical shift than the permutations of signs, because the GLM has to be retrained in each permutation. For our analysis we ran 119 permutations for this method, which means the uncertainty of the obtained *p*-values are slightly higher than for the other permutation method.

The minimum lag of 150 is part of an additional adjustment when performing the cyclical shifts, similar to the skipping done for the CV, as a measure against concerns raised by ***Harris*** (***2020a***) about the resulting discontinuities at the ends and at the seam of the shifted vector. When calculating the initial test statistic of the unshifted data, we exclude 75 data points in each end and 150 data points in the middle, and when calculating the test statistic of shifted data we exclude 75 data points on each side of the seam and 75 data points at each end, thus there are gaps of 150 data points around each discontinuity, similar to what we end up with during the skipping in the CV.

### Accounting for multiple comparisons when using permutation tests

In each step of the forward selection procedure we are considering multiple covariates to include. Conducting a hypothesis test only for the best performing covariate is a form of selective inference (***Taylor and Tibshirani, 2015***), a well known problem with few good and few general solutions. In a step of the forward selection procedure where *m* covariates are considered we are ignoring *m* − 1 hypothesis tests, meaning we essentially have a multiple testing problem (***Goeman and Solari, 2014***). As illustrated in Figure 11, this leads to an inflated type I error rate (we falsely reject true hypotheses more often than the set significance level *α*).

**Figure 11.**
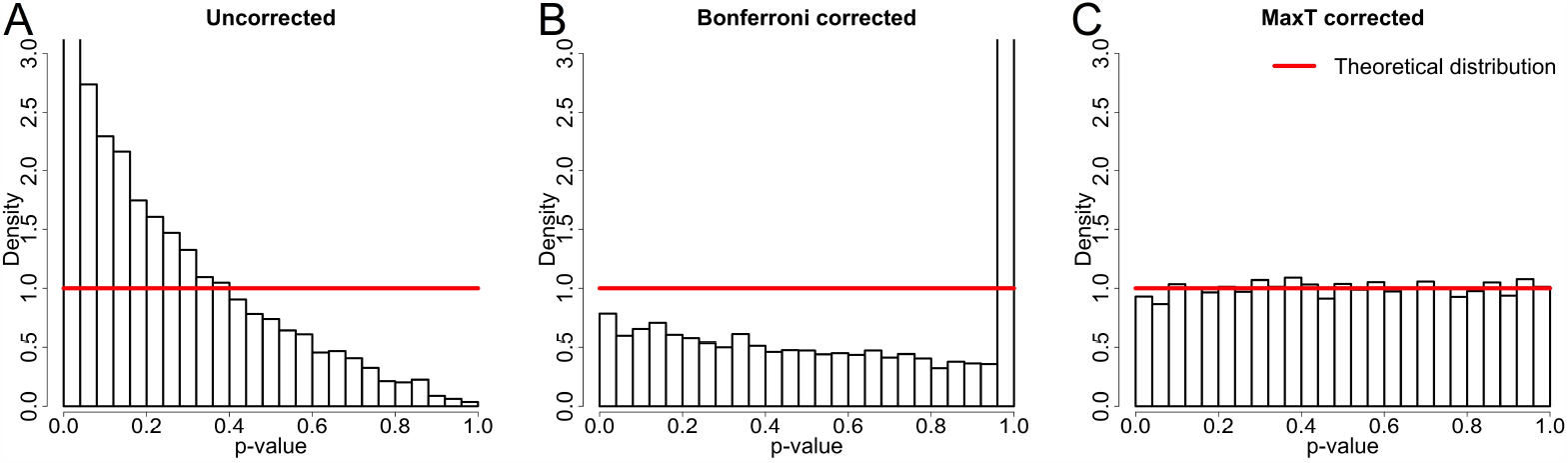
Illustration of how multiple comparisons results in too small *p*-values when not accounted for (**A**), and potential solutions using the Bonferroni (**B**) and maxT (**C**) correction methods.

We solve this issue by correcting our hypothesis tests to obtain familywise error rate (FWER) control in each step of the forward selection procedure. This means that we ensure at each step that the probability is at most *α* for rejecting any true hypothesis. A Bonferroni correction (***Goeman and Solari, 2014***) may be used and is always valid, although often conservative. For permutation testing we may however also use the maxT method by ***Westfall and Young*** (***1993***), making the adjustment not on the *p*-values, but by specifically using max *T*_*l*_ as the test statistic, where *T*_*l*_, *l* = 1, …, *m* are the individual test statistics for *m* covariates. Thus we create a shuffled distribution by calculating the maximum statistic over all candidate covariates in each permutation (meaning that it is not necessarily the same covariate that yields the maximum in each permutation), and comparing the observed maximum to this distribution.

For the sign-rank test method where we use sign flipping (mSR_MaxT_ and mSRR_MaxT_) we use the maxT correction, meaning our resulting test statistic is

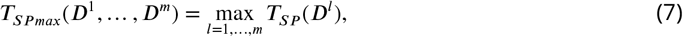

where each *D*^*l*^ is the vector containing the differences in cross-validated log-likelihood values between the current model and one of *m* candidate models (in our case *m* is 3 in the first step of the forward selection procedure, 2 in the second, and then 1 in the third, where only one candidate is tested and the corrected test statistic is just the single test statistic).

For the cyclical shift method we use the Bonferroni correction, because the in-sample log-likelihood that we use as a test statistic scales differently for covariates that are potentially parameterized with very different numbers of parameters. We also use the Bonferroni correction for the signed-rank test based method where we use the theoretical distribution to obtain the *p*-value (SR_Bonf_). In these cases the adjustment is made directly on the attained *p*-value, multiplying it by the number of candidates considered, *m*.

When using the Bonferroni correction it is important to keep in mind that when considering an increasing number of covariates, one needs to scale the number of permutations performed accordingly. If we consider 20 covariates, the correction is *p* ⋅ 20, meaning that to be able to obtain significant results with *α* = 0.05 we would need at least 399 permutations (as with *B* = 399 permutations and *m* = 20 covariates the smallest *p*-value we can obtain is 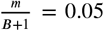). Ideally we would prefer even more permutations, to reduce the uncertainty of the estimated *p*-value. While this means the computational complexity can grow large, it also serves as an incentive to limit the number of covariates one tests simultaneously.

***Hardcastle et al. (2017***) limit the inflated familywise error rate by performing an additional hypothesis test at the end of the forward selection procedure, where the performance of the resulting model is compared to an empty one, concluding with no tuning if significance is not reached. In any given step there is however no correction, meaning the procedure may go on too far, and include enough irrelevant variables that the final test concludes that the final model should be discarded, even when there could be an intermediary model containing variables relevant to the neuron’s activity. Correction in each step is therefore preferable.

### Details on the simulated data

Here we describe the procedure we use to construct our simulated data. For our simulations we use *N* = 12000 data points. Covariates are simulated with temporal structure by first drawing *N* independent uniform variables from the interval (−2.5, 2.5), and then passing this vector through an exponential smoothing filter with scale parameter *β* = 20 (chosen to construct data appearing similar to the neural and behavioral data from ***Zong et al. (2022***), through visual inspection of the covariates themselves, and their autocorrelation function (see Figure 12)). Finally the covariates are bounded between −0.3 and 0.3 by replacing any exceeding value *v* > 0.3 or *v* < −0.3 with 0.3 − (*v* − 0.3) or −0.3 + (−0.3 − *v*), respectively, thus reflecting them back into the domain.

**Figure 12.**
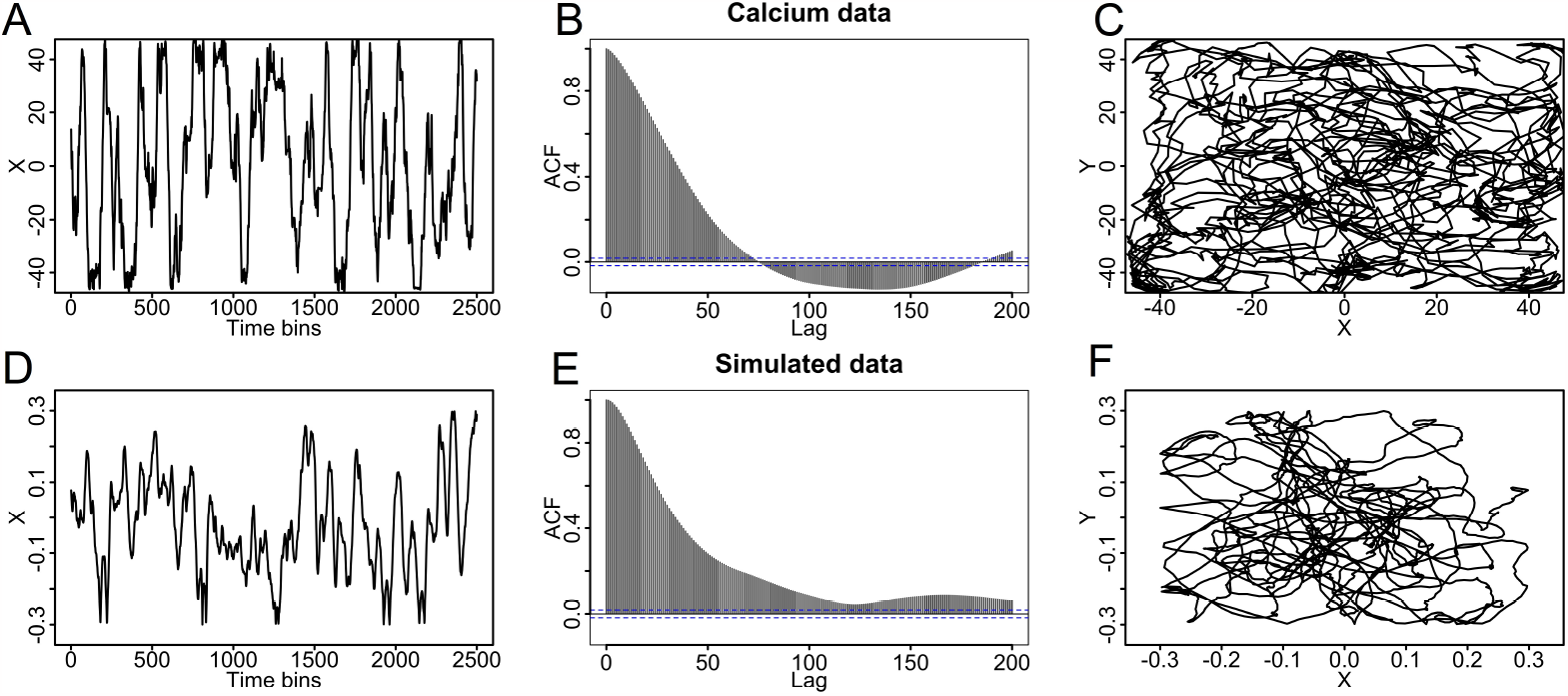
Comparison of real and simulated data. **A**: Animal X-position over time. **B**: Autocorrelation function of the animal’s X-position. **C**: X- and Y-position from the real mouse data. **D**: Simulated X-position. **E**: Autocorrelation function of the simulated X-position. **F**: Simulated X- and Y-position.

**Figure 13.**
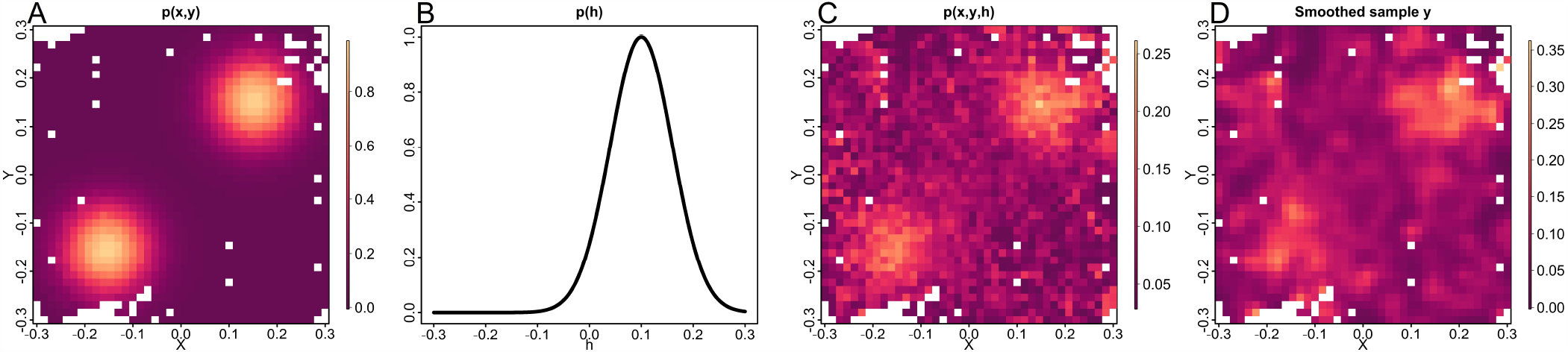
Illustration of how the event probability is constructed for the simulated data. **A**: The contribution X and Y have on the probability of an event, *p*_*xy*_. **B**: The contribution from *h, p*_*h*_. **C**: The resulting probability as a function of X and Y for one set of simulated (*x, y, h*). **D**: A smoothed ratemap of a sample of Bernoulli variables generated from *p*(*x, y, h*).

**Figure 14.**
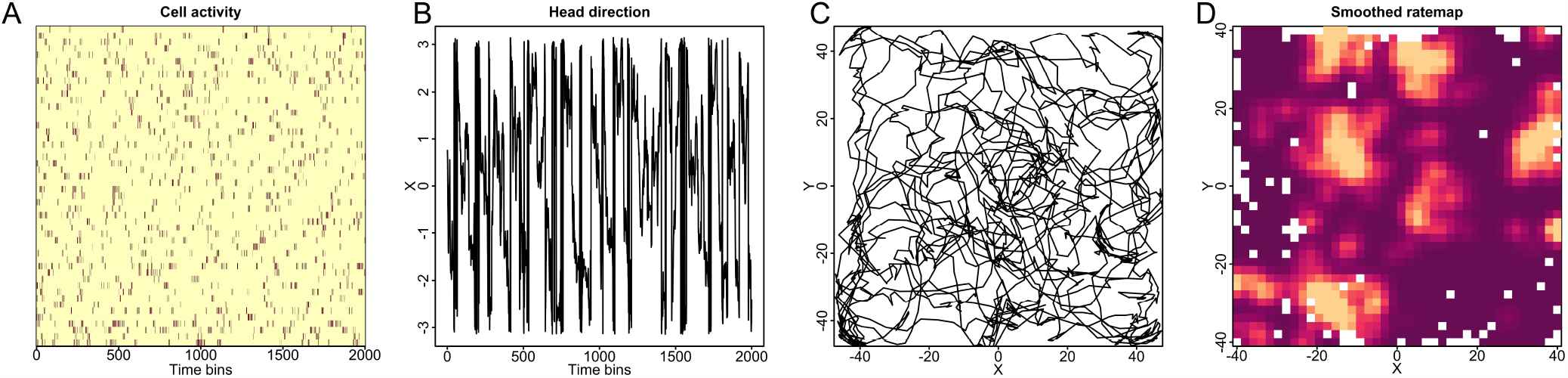
Illustration of calcium data. **A**: Raster of cell activity. **B**: Head direction of the animal over time. **C**: Animal position over time. **D**: Ratemap of the cell activity over position for a single neuron.

In a single simulation we generate 5 such covariates independently of each other. One of these, *h*, is treated as unobserved with respect to the covariate selection process, but is used to generate the response variable. The next two are treated as observed, one-dimensional covariates, while the last two are combined and treated as one two-dimensional covariate with *x*- and *y*-coordinates. When fitting the GLM we represent the covariates using natural cubic splines, using the R packages splines. We use 5 evenly spaced internal knots for the one-dimensional covariates and 2 evenly spaced internal knots for each dimension of the two-dimensional variable.

We generate two sets of simulated data, one where only *h* affects the underlying probability of firing, and one where the two-dimensional covariate also affects the probability. We generate new covariates for each simulated response variable (cell activity). The probability in each bin is first modelled using Gaussian bumps plus a base probability. The contribution from *h* is

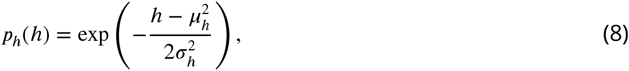

where the center of the bump is at *μ*_*h*_ = 0.1 and the standard deviation is *σ*_*h*_ = 0.06. For the data where the position (*x, y*) is relevant, the contribution, constructed using two two-dimensional Gaussian bumps, is

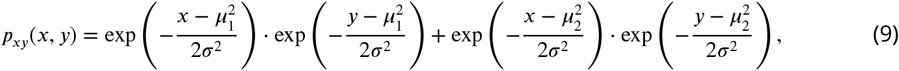

where we have centers at (*μ*_1_, *μ*_1_) = (−0.15, −0.15) and (*μ*_2_, *μ*_2_) = (0.15, 0.15) and standard deviation *σ* = 0.06. The combined probability is then

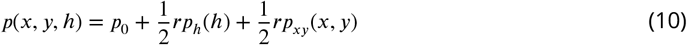

when position is relevant, and

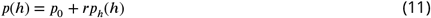

otherwise, where *p*_0_ = 0.03 is the base probability, and *r* is a scaling parameter set to 0.25 for the main analysis, in order to yield an overall rate of events similar to the real data. The parameters are set by trial and error in an attempt to construct ratemaps similar to what one could observe with actual neural data, albeit slightly less complex than the grid cells observed in the real data we explore in order to reduce computational labor. By increasing *r* we can make the response depend more strongly on the hidden covariate and on position, if it is included. With *r* = 1 all the methods compared in Results on simulated data will correctly classify the cells as position-tuned in essentially every trial (meaning we cannot say anything meaningful about their differences in test power).

### Details on the calcium data

The data used for the analysis of calcium data is from ***Zong et al. (2022***) and ***Obenhaus et al. (2022***), where they use a 2-photon miniscope to obtain calcium images of freely moving mice. We are considering specifically the dataset from mouse 97045, recorded March 17th, 2021, with calcium imaging of the medial entorhinal cortex. The neural activity is sampled at 7.5Hz and deconvolved using Suite2P (***Pachitariu et al., 2016***). We binarize their filtered events (setting any non-zero value to 1) and treat these as Bernoulli variables. They analyze the tracking data, which is sampled at 15Hz, using DeepLabCut (***Mathis et al., 2018***), and combine the tracking and neural data using their own software, NATEX. For our analyses we have used the 7.5Hz rate, combining every two time frames of the tracking data by taking the average.

The session we have used lasts for one hour, of which we use 12000 time bins (26 minutes and 40 seconds) of the first half as our correctly matched data, and the neural data from the 12000 first time bins of the second half to create our mismatched data when combined with the tracking of the first part. The session used has a total of 488 neurons, of which we consider the 249 neurons that have an event in at least 2% of the time bins in each of our two 12000 time bin parts. 147 of these are classified by ***Zong et al. (2022***) as grid cells.

When fitting the GLM we represent the covariates using natural cubic splines. For head direction we use periodic B-splines, with the R package pbs. For head direction we use 6 evenly spaced internal knots and for speed we use 5, which results in the same number of parameters since the former is periodic. For position we use 4 evenly spaced internal knots for each dimension.

## Acknowledgments

We would like to thank Ingeborg Hem and Mette Langaas for their comments on the manuscript, and thank the Department of Mathematical Sciences (NTNU). We would also like to thank ***Zong et al. (2022***) and ***Obenhaus et al. (2022***) for making their data publicly available. This work was supported by a grant by the Research Council of Norway (iMOD, NFR grant #325114).

